# Genomic connectivity and adaptation signals of the freshwater sponge *Ephydatia muelleri* across its distribution

**DOI:** 10.1101/2025.01.21.634144

**Authors:** Robert Cassidy, Laura de la Cruz, Konstantina Mitsi, Carles Galià-Camps, Ana Benítez-López, Carlota Gracia-Sancha, Jose Lorente-Sorolla, Almudena Álvarez, Rocío Mozo, Stephen Kolomyjec, Scott Nichols, Renata Manconi, Raquel Pereira, Karen Evans, Valeria Itskovitch, April L. Horton, Sally P. Leys, Sergio Taboada, Ana Riesgo

## Abstract

Freshwater sponges fulfill critical ecological functions, including maintaining water quality, regulating nutrient dynamics, offering habitats for diverse taxa, and serving as a vital food source for various species. However, their patterns of dispersal and genetic connectivity remain inadequately understood, posing significant challenges to effective conservation assessments. We examined genetic connectivity and genetic adaptation to local environmental conditions in *Ephydatia muelleri* across its geographic range using ddRADseq-derived SNPs from 106 individuals collected from 11 localities spanning North America, Europe, and Asia. Analysis of 3,182 neutral SNPs revealed low connectivity and strong genetic structure among regions within two main genetic clusters of North America and Eurasia, while 115 SNPs identified to be under selection showed considerable evidence for differentiated, polygenic adaptation to light and temperature conditions across sampled locations, as well as selection on gene regulatory processes. These findings align with the “monopolization hypothesis”, suggesting that historical climatic and geological conditions of the Last Glacial Maximum, including habitat expansion, contraction, and natural barriers, have contributed more to the current genetic structure of *E. muelleri* populations than contemporary gene flow, which is restricted by monopolistic habitat colonization by this species. Our results provide novel support for ecological theory on dispersal in aquatic invertebrates, as well as insights into the plasticity of *E. muelleri* in the face of varying environmental conditions that are fundamentally important for freshwater ecosystem conservation.

## 2. Introduction

From microscopic plankton to enormous aquatic mammals, freshwater systems—such as rivers, lakes, marshes, and streams—support a diverse array of life, whose dynamics are vital for maintaining the health and balance of ecosystems, yet they are among the most strongly altered ecosystems with an overproportional loss of biodiversity (Geist 2011). The most important threats to freshwater ecosystems include habitat loss and alteration, direct mortality from exploitation, invasive species, emerging infectious disease, pollution and climate change, threatening their equilibrium by altering their communities with cascading ecosystem effects (Geist 2011). Wildlife population threats frequently work in concert, and genetic factors play a major role in the challenges wildlife faces. For example, inbreeding and genetic diversity loss, brought on by population fragmentation and reductions, can both directly lower population fitness and a population’s capacity to adapt to new environmental conditions (Pauls et al. 2014; Hohenlohe et al. 2021). In this sense, measuring and reducing threats to wildlife populations requires an understanding of genetic and genomic features and how they interact with environmental factors in nature to effectively design conservation strategies (Pauls et al. 2014; Hohenlohe et al. 2021).

By offering previously unprecedented insight into the histories of thousands of loci, population genomics is transforming our understanding of evolutionary processes in natural populations and revealing the effects of migration, genetic drift, natural selection, and mutation on the spatial distribution of species (Hohenlohe et al., 2021; Luikart et al., 2019). From methods like Double Digest Restriction-Site Associated DNA Sequencing (ddRADseq), which can produce thousands of single nucleotide polymorphisms (SNPs), to whole genome sequencing approaches, the field has progressively moved from genotyping to genome typing (Luikart et al., 2003). The next limitation, related to the availability of chromosome-scale genomes for the target species, is being addressed by the large-scale production of quality genomes for a great variety of organisms. This enables the study of adaptation patterns by scanning for outlier loci, which can reveal regions likely subjected to strong diversifying or stabilizing natural selection, offering valuable insights for conservation efforts (Formenti et al., 2022; Theissinger et al., 2023). In this context of biodiversity conservation, most efforts have focused on vertebrates (Donaldson et al., 2016), often overlooking the importance of invertebrates, especially in aquatic systems. These species may experience unique environmental and anthropogenic pressures, yet we still lack a clear understanding of their resilience to such challenges (Collier et al., 2016).

Sponges are among the most common and abundant invertebrates in aquatic ecosystems, with 99% of them inhabiting the marine realm (Van Soest et al., 2012). However, only the order Spongillida has been present in freshwater systems since the Permo-Carboniferous (about 300 Mya; (Pronzato et al., 2017)). Spongillids have a global distribution and diversified significantly during the Paleogene, allowing adaptation to diverse climates (Manconi & Pronzato, 2016a; Pisera et al., 2016; Pronzato et al., 2017). Spongillida currently includes 7 families, 47 genera, and 236 species worldwide, with high endemism except for certain widespread genera like *Spongilla* and *Ephydatia* (Manconi & Pronzato, 2016b). While cosmopolitanism is nearly nonexistent in marine sponges, freshwater species have exceptionally large distribution ranges. In particular, *Ephydatia muelleri* is widely distributed across temperate regions of the Holarctic, including Europe, North America, and Asia (Manconi & Pronzato, 2016b). To disperse over such large distribution ranges, freshwater sponges rely on specialized dormant gemmules, resilient structures with totipotent cells encased in a protective layer (Maldonado & Riesgo, 2009; Manconi & Pronzato, 2016b; Ungemach et al., 1997). These gemmules can survive adverse conditions in a cryptobiotic state, reactivating to restore functions such as filtration and reproduction when conditions improve (Kenny et al., 2020; Manconi & Pronzato, 2016a). Gemmules facilitate dispersal across fragmented habitats through both biotic and abiotic vectors, such as mammals, birds, streams, and wind. Structural adaptations, including spiny spicules and a spongin-based pneumatic layer, enhance attachment to animal fur or feathers, facilitating efficient transport (Maldonado & Riesgo, 2009; Pronzato & Manconi, 1994) but also providing buoyancy to ensure drifting.

Freshwater sponges face a range of ecological challenges in their habitats, including fluctuating environmental conditions, habitat fragmentation, and varied resource availability (Pronzato & Manconi, 1994). To survive in these dynamic environments, they have adaptively acquired a variety of physiological, morphological, and reproductive strategies. These mechanisms include asexual reproduction through fragmentation, where individual sponges can break into pieces that regenerate into new individuals (Frost, 1991; Manconi & Pronzato, 2002), and physiological adaptations such as the ability to adjust filtration rates or alter growth patterns in response to changing conditions (Cartwright et al., 2024; Francis et al., 1982). These adaptations are crucial for ensuring that sponges can withstand environmental stressors and thrive in diverse habitats, whether they are experiencing fluctuating water quality, temperature changes, or varying nutrient availability. Together, both adaptation and dispersal mechanisms significantly influence the genetic structure of freshwater sponge populations. The adaptive strategies that allow sponges to survive in specific environmental conditions shape the genetic makeup of local populations, leading to the accumulation of unique genetic signatures (Lucentini et al., 2013). Meanwhile, the dispersal of gemmules facilitates gene flow between populations, preventing genetic isolation and promoting genetic diversity across broader geographical areas (Li et al., 2018). These interconnected processes—adaptation to local conditions and dispersal across fragmented habitats—ultimately determine the evolutionary trajectory of freshwater sponge populations, influencing their genetic structure and dynamics over time. Despite extensive research on the population structure of various freshwater invertebrates, which show low genetic diversity and isolation-by-distance (Bohonak & Jenkins, 2003; Garrison et al., 2021), our understanding of population structure in freshwater sponges remains limited. Available studies suggest strong genetic differentiation in spongillid populations, driven by isolation, physical barriers, and the low dispersal capacities of their larvae (Li et al., 2018; Lucentini et al., 2013; Yakhnenko et al., 2021, 2024). However, these studies predominantly rely on traditional markers and microsatellites, which are less effective than single nucleotide polymorphisms (SNPs) for detecting subtle connectivity patterns (Andrews et al., 2016). Notably, SNPs have already demonstrated significant potential in sponge research (Leiva et al., 2019; Taboada et al., 2022, 2023).

Our study adopts a genomic approach to explore the genetic connectivity and adaptive patterns of the freshwater sponge *Ephydatia muelleri* across its range, with the goal of elucidating its evolutionary history, current population structure, and adaptive mechanisms, thereby providing insights to inform conservation strategies. *Ephydatia muelleri*, a freshwater demosponge widespread in temperate regions across the northern hemisphere, including parts of Europe, North America, and Iceland (Cardenas et al., 2012; Hall et al., 2021). It reproduces both sexually and asexually, with a sexual reproductive period from November to July, when it releases larvae that settle on suitable substrates (Mukai, 1990; Sallin, 1995). Asexual reproduction occurs through fragmentation and gemmulation, with hardy gemmules that withstand extreme conditions like drought and anoxia, facilitating the species’ survival, dispersal, and colonization of diverse aquatic habitats (Frost, 1991; Manconi & Pronzato, 2002; Sallin, 1995). With its broad geographic range, asexual reproduction strategies (Frost, 1991; Sallin, 1995), and well-established precedents in natural and laboratory studies (Kenny et al., 2020) *Ephydatia muelleri* serves as an ideal model for examining genetic variability and evolutionary dynamics across time and space. For this purpose, we employ a ddRADseq genomic approach on 106 specimens across its whole distribution range to evaluate *E. muelleri* population structure, coancestry patterns, migration rates among localities, and ancient demographic patterns, as well as environmental-linked loci which could respond to local adaptation and their potential biological function.

## 3. Materials and Methods

### 3.1. Sample Collection and Preservation

A total of 106 specimens of *Ephydatia muelleri* were collected from 11 sites across the species’ distribution range. These sites included O’Connor Lake (Canada), various rivers in Canada, Colorado, Michigan, Maine, Ireland, Spain, the United Kingdom, Sardinia, Sweden, Estonia, and Lake Baikal (Table 1; Figure 1). Samples were gathered manually from lakes, rivers, and reservoirs at depths up to 40 cm, and preliminarily identified based on morphology of spicules. Specimens were preserved in 96% ethanol, DNA/RNA shield reagent (Zymo Research) or RNAlater (Invitrogen) and stored at −20°C for further analysis.

**Table 1.** Population genetic statistics for *E. muelleri*. Samples grouped in eight areas and at the global level. N: number of samples, *He*: expected heterozygosity, *Ho*: observed heterozygosity.

| Sampling group | N | Private alleles | Private alleles/N | $H_e$ | $H_o$ |
| --- | --- | --- | --- | --- | --- |
| O'Connor lake (Canada) | 10 | 84 | 6,46 | $0.044 \pm 0.002$ | $0.047 \pm 0.003$ |
| Nanaimo & Sooke river (Canada) | 13 | 132 | 13,2 | $0.075 \pm 0.002$ | $0.063 \pm 0.003$ |
| Colorado | 4 | 29 | 7,25 | $0.043 \pm 0.002$ | $0.080 \pm 0.005$ |
| Michigan | 21 | 555 | 26,42 | $0.105 \pm 0.003$ | $0.076 \pm 0.002$ |
| Maine | 13 | 268 | 20,61 | $0.087 \pm 0.003$ | $0.071 \pm 0.003$ |
| Spain | 30 | 52 | 1,73 | $0.018 \pm 0.001$ | $0.027 \pm 0.003$ |
| Europe without Spain | 10 | 409 | 40,9 | $0.084 \pm 0.003$ | $0.058 \pm 0.002$ |
| Baikal | 5 | 139 | 27,8 | $0.058 \pm 0.003$ | $0.062 \pm 0.003$ |
| **Total** | 106 | - | - | $0.108 \pm 0.002$ | $0.050 \pm 0.001$ |

**Figure 1.**
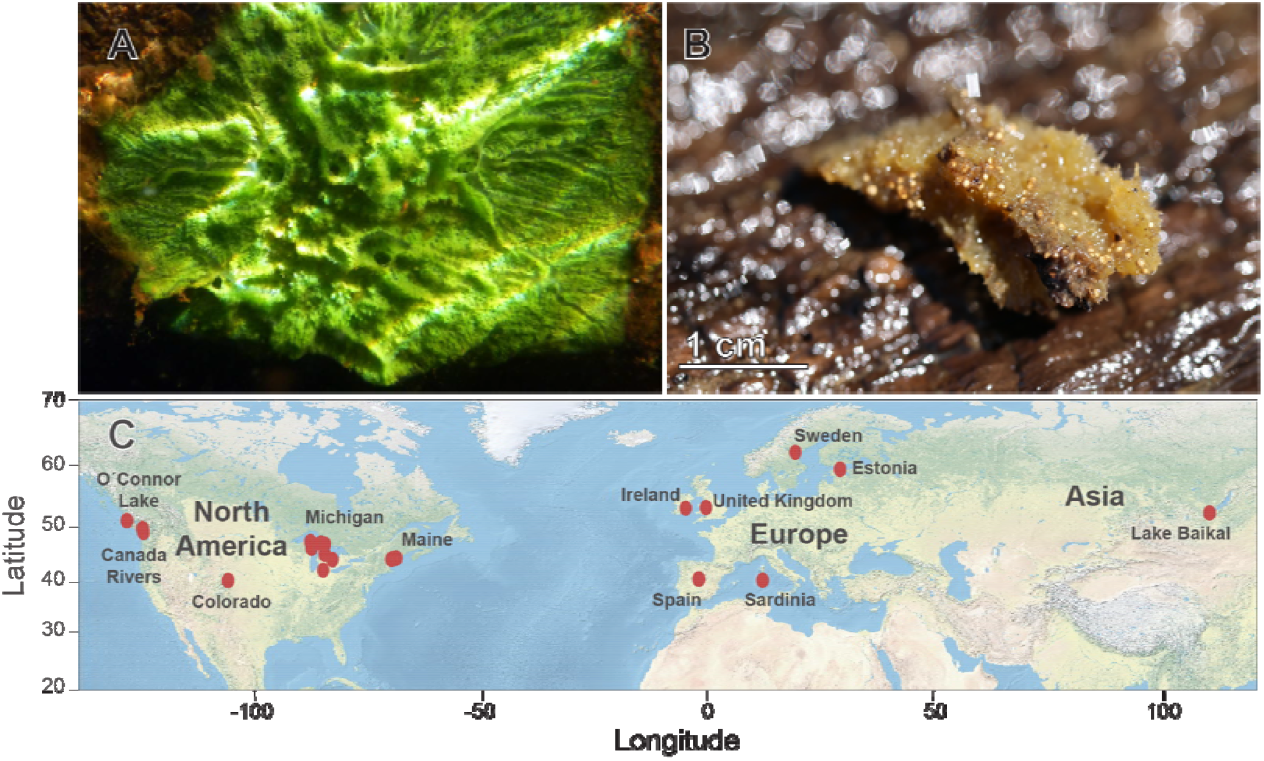
*Ephydatia muelleri* from **A**. Sooke River, Canada (picture by Sally Leys) and **B**. Embalse de los Morales reservoir (Spain), displaying gemmules within the tissue (picture by Ana Ramón-Laca). **C**. Map with locations of the sampled sites.

### 3.2. DNA Extraction, Barcoding and Haplotype networks

Due to the high phenotypic plasticity of freshwater sponges, DNA barcoding was performed on 98 specimens to confirm their identification as the target species. Genomic DNA was extracted from approximately 1 cm^3^of each sample using the Speedtools Tissue DNA extraction kit. DNA concentration was measured with the Qubit 1x dsDNA High Sensitivity assay kit (Thermo Fisher), and quality was assessed using a NanoDrop spectrophotometer (Thermo Fisher). The Internal Transcribed Spacer (ITS1) region was amplified using the primers FS-ITSF 5’-TACACACCGCCCGTCGCTACTA-3’ and FS-ITS-R 5’-CTYYGACGTGCCTTTCCAGGT-3’ (Itskovich et al., 2008). The PCR amplification protocol consisted of an initial activation cycle at 94°C for 2 minutes, followed by 40 cycles of denaturation at 94°C for 60 seconds, annealing at 55°C for 60 seconds, elongation at 72°C for 60 seconds, and a final extension at 72°C for 8 minutes. Each PCR reaction was conducted in a 25 _μ_L volume, which included 12.5 _μ_L of 2x PCRBIO Taq Mix Red (PCR Biosystems, UK), 1 _μ_L of each primer (10 _μ_M), and 1 _μ_L of DNA. Successful amplification was verified by electrophoresis on a 1.5% agarose gel at 90 V for 45 minutes. The amplified samples were then sequenced using Sanger sequencing with the same primers at the Macrogen laboratory (Spain). The resulting sequences were compared against known databases using the BLAST program to confirm species identification based on significant matches with ITS sequences in the NCBI repository. Then, sequences of ITS were aligned with MAFFT (Katoh et al. 2002) and implemented in PopArt (Leigh et al., 2015) for the construction of TCS haplotype networks.

### 3.3. ddRADseq Library Preparation and Sequencing

ddRADseq genomic libraries were prepared for all samples following the protocol of Peterson et al. (2012) with modifications according to Combosch et al. (2017). Double-stranded genomic DNA (500 ng) was digested using high-fidelity restriction enzymes EcoRI and BfaI (New England Biolabs) for 6 hours at 37°C. The resulting digested fragments were cleaned by manual pipetting with Agencourt AMPure beads (1.5x volume ratio; Beckham Coulter) and quantified using a Qubit dsDNA HS assay. These fragments were then ligated to custom P1 and P2 adapters, which included unique barcodes for each sample and hybridization sites for the primers. After adapter ligation, the samples were pooled into libraries, with a maximum of 12 individuals per library. The libraries were cleaned again using AMPure beads (1.5x volume ratio) and size-selected (200–400 bp) using a Blue Pippin Prep system (Sage Science). Each library was amplified by PCR using Phusion polymerase (Thermo Scientific) and library-specific PCR primers to allow multiplexing of the libraries. The PCR program used was 98°C for 30 seconds, followed by 12 cycles of 98°C for 10 seconds, 65°C for 30 seconds, and 72°C for 90 seconds, with a final extension at 72°C for 10 minutes. The resulting PCR products were cleaned again using AMPure beads (1.5x volume ratio), quantified with a Qubit dsDNA HS assay, and the fragment size pattern was checked on a Tapestation 2200 (Agilent Technologies). Finally, the library concentrations were normalized and pooled for sequencing in a single run on an Illumina NovaSeq 6000 (2 × 150 bp) at Novogene Europe (Cambridge, UK).

### 3.4. Assembly and Filtering of ddRADseq Loci

Following sequencing, quality filtering and loci assembly were conducted using Stacks v2.57 (Catchen et al., 2013). The “process_radtags” module was used to filter raw reads by removing low-quality sequences, those missing barcodes, or with incomplete restriction sites. RAD-tags with minimal divergence, such as adapter mismatches (allowing up to two mismatches with the parameter --adapter_mm = 2), were retained using the “-r” parameter. Reads were trimmed to 145 bp with the “-t” parameter to enhance SNP detection accuracy. After filtering, 43,651,303 sequences were retained from an initial 44,103,246, averaging 4,270,730 sequences per individual (range: 206,006 to 30,804,006).

High-quality sequences were aligned to the *E. muelleri* reference genome (Kenny, 2020) using Bowtie2. A reference genome index was first constructed to enable efficient sequence searches. Alignments were conducted with paired-end sequence files specified via the “-1” and “-2” parameters, along with the “--very-sensitive-local” setting to enhance alignment sensitivity, accommodating partial matches when reads diverged from the reference. This approach ensured accurate alignments, even for reads with significant variations. A bash command was used to extract alignment percentages from output files, averaging 80% for the dataset, indicating high coverage and optimized alignment. Following alignment, the Stacks tool “ref_map.pl” was employed to process the aligned data. A population map (“--popmap”) and sample directory (“--samples”) were provided, and “gstacks” was run to assemble loci and identify SNPs based on read alignment positions.

The “populations” tool in Stacks was used to extract SNPs present in at least 80% of individuals (r = 0.8). This threshold was chosen because 0.8 is an appropriate value that strikes a balance between being neither overly restrictive nor too broad, allowing for a representative selection of SNPs without excluding valuable data or including too many non-informative variants. To reduce linkage disequilibrium, the first SNP from each RAD-tag was selected with the “--write_single_SNP” parameter. Following Roesti et al. (2012), only SNPs with a minor allele frequency (--min_maf) above 0.05 were retained. SNPs failing Hardy-Weinberg equilibrium (p ≤ 0.05) in at least two areas or showing excess heterozygosity (Ho > 0.5) were removed, per Hohenlohe et al. (2011). Further filtering was performed using PLINK v1.9 (Purcell et al., 2007) with the “--indep-pairwise 50 10 0.5” command. Note that these filters were not applied to run fineRADstructure as linked regions may capture insightful information regarding coancestry. This step removed highly correlated SNPs by recalculating linkage disequilibrium across 50-SNP windows every 10 SNPs, ensuring uncorrelated data for downstream analyses. The final SNP matrix was visualized with the Matrix Condenser interface, revealing global missing data of 10.05%, ranging from 4.39% to 18.29% per individual. These values supported retaining all 106 individuals for further analyses.

### 3.5. Population Genomics Analysis

To assess neutral population structure and connectivity, SNPs identified as being under selection through a combination of Arlequin, BayeScan, Baypass, LFMM, and pRDA analyses (described in more detail in section 3.8) were filtered out, and only the resulting 3,182 neutral, filtered SNPs were used for subsequent analyses.

Population genetic statistics were computed from the neutral dataset. First, samples were grouped into eight regions: O’Connor Lake (Canada), Nanaimo and Sooke River (Canada), Colorado, the Great Lakes (Michigan), Maine, Spain, Europe excluding Spain (comprising the United Kingdom, Ireland, Estonia, Sardinia, and Sweden), and Asia (Lake Baikal). This grouping was selected not only to reduce the initial 11 sites due to low and uneven sample sizes in certain locations, thereby ensuring more robust parameter estimations, but also in response to the absence of preexisting data on the population genetics of freshwater sponges across such a broad habitat range. In this context, we performed an a priori Discriminant Analysis of Principal Components (DAPC) with less stringent criteria to guide the delineation of populations for subsequent analyses. These preliminary analyses demonstrated that population differentiation could be enhanced by effectively separating lake and river populations in Canada and distinguishing populations in Spain from those in the rest of Europe. Genetic metrics, including expected heterozygosity (He) and private alleles, were calculated using Stacks. He, a measure of genetic diversity, quantifies the probability that two randomly selected alleles from a population are different. Private alleles, which are unique to specific populations, were also assessed to further understand population differentiation.

Population structure was evaluated using three methods: STRUCTURE v2.3 (Pritchard et al., 2000), Discriminant Analysis of Principal Components (DAPC) from the adegenet v2.1.10 package in R (Jombart et al., 2010), and fineRADstructure v1.0.2 (Malinsky et al., 2018). These three approaches utilized the previously explained groupings. STRUCTURE was run with 200,000 MCMC iterations under the admixture model, with a burn-in of 100,000 iterations, evaluating K values from 1 to 8 with 15 replicates for each K value. CLUMPP version 1.1.2 (Jakobsson and Rosenberg, 2007) was used to identify the most probable number of clusters and to average the ancestry proportions of each individual across K replicates.

Population structure was analyzed using Discriminant Analysis of Principal Components (DAPC) with the *snapclust* function in the *adegenet* package in R. Optimal genetic clusters were determined using Akaike (AIC), Bayesian (BIC), and Kullback (KIC) information criteria combined with the k-means algorithm, allowing up to 16 clusters and 100 iterations. The cluster number with the lowest AIC, BIC, and KIC values was selected. The optimal number of principal components (PCs) was identified using the *xvalDapc* validation function with 1,000 replicates, minimizing the mean squared error (MSE) while retaining maximum variance. Individual assignment probabilities and cluster scatterplots were visualized using *assignplot* and *scatter*.*plot*, respectively.

To assess shared ancestry among the analyzed individuals of *E. muelleri* and to support the STRUCTURE and DAPC analyses, the fineRADstructure program was also used. This program leverages the information from all SNPs present in RAD-tags to provide high-resolution results on coancestry between individuals. The analysis was conducted using all individuals with 43,000 SNPs, obtained by running “populations” in Stacks with 3,339 neutral SNPs, allowing the inclusion of SNPs in linkage disequilibrium by deselecting the “--write_single_SNP” option and skipping preliminary filtering with PLINK. The fineRADstructure analysis was run with default values: 100,000 for -x, 100,000 for -y, and 1,000 for -z, assigning individuals to populations, and 10,000 for -x in tree construction. The results were graphically interpreted using the “Finestructure R Library” and the fineRADstructurePlot.R script, both provided in the fineRADstructure package for R.

In addition to the genetic structure analyses described above, a molecular variance analysis (AMOVA) was also performed using Arlequin to evaluate the significance of differentiation between the samples from the East (Eurasia) and the West (North America). The Eastern samples included Spain, Europe (United Kingdom, Ireland, Estonia, Sardinia, Sweden), and Lake Baikal, while the Western samples included O’Connor Lake (Canada), Nanaimo and Sooke River (Canada), Colorado, the Great Lakes (Michigan), and Maine. The AMOVA analysis was conducted using the Standard AMOVA option, with the missing data level per locus set at 1 and 20,000 permutations, resulting in 3,182 usable loci. The samples were grouped into a total of 11 populations to increase the statistical power of the comparisons in the analysis.

Finally, pairwise F_ST_ values were estimated to measure genetic differentiation between the eight sample groupings using Arlequin. The analysis was configured to allow up to 1% missing data per locus, and 20,000 permutations were performed. F_ST_ values range from 0 (no differentiation between the compared populations) to 1 (maximum differentiation between the compared populations).

### 3.6. Demography and Migration Patterns

The neutral SNPs dataset was transformed into folded Site Frequency Spectrum (SFS) files with the software easySFS.py (https://github.com/isaacovercast/easySFS) for demographic purposes. We used the neutral SNPs dataset as adaptive loci might generate biases in the resulting demographic history. We modeled the historical demographic events with the software Stairwayplot2 (Liu & Fu., 2020). Recent studies using RADseq data with this software have shown that drastic changes in effective population size can mirror colonization and bottleneck processes (Galià-Camps et al., 2024). We used a single SFS file including all the individuals, regardless of the locality or genetic cluster, and kept 158 haplotypes to generate the species SFS file. We established a generation time of 1 year since most of continental sponge species have an annual cycle (Gugel, 2001), and a mutation rate per site and generation of 1×10e-9 as spongillids are known to have low mutation rates in comparison to other sponge taxa (Maas et al., 2023). We ignored singletons as recommended by the software developer (Liu & Fu., 2020), generated 200 input files to estimate the demographic patterns, and visualized the results with ggplot2.

To identify current patterns of gene flow in the study area, Nei’s G_ST_ method was used to estimate contemporary migration between geographic areas, employing the divMigrate function from the R package diverRsity v1.9.90 (Keenan et al., 2013). Additionally, a filtering threshold of 0.3 was selected for the analysis, meaning that only gene flows with values above this threshold were considered. This value was chosen to ensure that detected migrations were significant and reflected relevant genetic connections between the studied geographic areas.

### 3.7. Detection of putative SNPs under selection and functional assignment

In order to examine possible mechanisms of local adaptation in populations of *E. muelleri*, we analyzed our SNP datasets to detect signatures of natural selection across sampled individuals’ reduced genomes. Different types of natural selection (e.g., monogenic vs polygenic) can produce diverse signatures of selection, and these signatures can be further confounded by nuances of population structure and gene flow (de Villemereuil et al., 2014; Rellstab et al., 2015; Narum & Hess, 2011; Excoffier et al., 2009). Existing methods of detecting adaptive assesses relationships can be broadly categorized into Gene-Environment Association (GEA) methods, which assess association between genetic data and relevant environmental variables, and Genome Scan (GS) methods, which assess statistical differentiation along the genome according to mathematical models without consideration for environmental associations. Among other considerations, some GEA analyses, such as RDAs, have been shown to better account for weak, polygenic signatures of selection, while GS methods can be an effective method of detecting selection for large-effect loci or when the impact of environmental variables is unknown (Lotterhos & Whitlock, 2015; Dallaire et al., 2021). Given the lack of consensus regarding the best method, and in order to assess a wide range of potential adaptive loci, we employed five different methods, including both GS and GEA approaches, to detect adaptive SNPs. GS techniques used included BayeScan (Foll and Gaggiotti, 2008), Arlequin (Excoffier and Lischer, 2010), and Baypass v2.4 (Gautier, 2015), while genome-environment association methods included LFMM and pRDA analyses using the *vegan* v2.6-8 (Oksanen et al., 2022) and *lfmm* v1.1 (Caye et al., 2019) packages in R, respectively.

For Arlequin, which detects outlier SNPs based on F-statistics, the missing data threshold per locus was set to 0.05, and a non-hierarchical island model, which assumes that populations are partially isolated from each other like islands, was used. Then, 100,000 simulations and 100 demonstrations per group were performed, and the resulting p-values were adjusted using the “p.adjust” function in R with the “BH” method by Benjamini and Hochberg (1995) to control the false discovery rate (FDR) and generate q-values for putative SNPs. BayeScan, which follows a Bayesian approach using allele frequency differences to identify SNPs under selection based on a multinomial-Dirichlet model, was then run according to default parameters, and SNPs with q-value > 0.05 were considered outliers for both methods. Baypass, which incorporates XtX statistics and allele frequency differentiation to calculate outlier SNPs, was run first on the LD-pruned SNP dataset with default parameters in order to create an omega matrix accounting for linkages and population structure. Because Baypass outputs can vary when runs are repeated, the resulting omega matrix was then applied in five consecutive runs of the Baypass program on a SNP dataset that had been filtered for HWE, Ho, and minimum allele frequency according to previously described thresholds, but had not been LD pruned. Of these five runs, the output with the median average XtX value was selected for analysis. Following the recommendations of Gautier (2015), XtX thresholds were then calculated by simulating a neutral SNP distribution and running Baypass on the simulated genotypes and were then applied as FDR cut-offs to identify outlier SNPs in the real dataset (FDR < 0.5).

For GEA methods, a pRDA and LFMM approaches were chosen for their ability to account for population structure and identify associations between SNPs and environmental variables. Freshwater environmental variables with potential biological significance for *E. muelleri* were selected from the Copernicus DynQual (Jones et al. 2023) and GeoFresh (Domisch et al. 2024) databases. We extracted 49 environmental variables related to water quality and temperature of the sampled site, and topography, soil characteristics and land cover variables at the subcatchment level (see details in Appendix S1) from our sampling sites using the R packages sf v1.0-19 (Pebesma, 2018), raster v3.6-30 (Hijmans 2024), and ncdf4 v1.23 (Pierce 2024). We assessed multicollinearity between variables with the *psych* package v2.4.6 in R (Revelle & Revelle, 2015) and filtered out collinear variables (Pearson’s rho > 0.6), resulting in a total of 14 environmental variables. Then, the same set of filtered, linked SNPs used for Baypass analysis was loaded and missing data was imputed with the most common genotype data for each SNP in the dataset.

Next, a PCA was performed on the SNP dataset filtered for linkage disequilibrium and without potential outlier SNPs identified by genome scan methods in order to best characterize the neutral structure of the sample populations (axes = 3). This structure was run as a condition in a pRDA with the scale-transformed environmental factors using the *rda* function in *vegan*. Then, ANOVA, VIF and R^2^ analyses were run, and outlier SNPs were calculated based with z = 3.

For LFMM analysis, in order to maintain an optimal genome inflation factor (GIF) profile, the dataset of LD-pruned SNPs was used. A PCA was performed using on the SNP data and PC1 and PC2 were selected for respective LFMM analyses because they explained similar levels of variance. Seven latent factors (K = 7) were assigned according to the populations identified in the DAPC and STRUCTURE analyses, and the *lfmm_ridge* and *lfmm_test* functions of the *lfmm* package were used to run latent factor mixed models for each PC. p-values were then converted to q-values according to FDR procedure and using the *qvalue* package by Benjamini and Hochberg (2005), with q-values < 0.05 considered to be significant.

In order to determine possible functions of putative SNPs under selection, candidate loci identified through the aforementioned combination of models were run aligned against the loci generated by Stacks using the BLAST program’s *blastn* function with e-value 0.005 as threshold (Altschul et al. 1997), and the resulting gene identities were implemented in *shinyGO* in order to obtain enriched gene ontologies, using the human genome for comparison and the *Ephydatia muelleri* genome as background (Ge et al. 2020). Then, GO terms were run in REVIGO to visualize categorizations using treemaps (Supek et al., 2011).

## 4. Results

### 4.1. Population structure and connectivity using ITS and neutral SNPs

We obtained four main ITS haplotypes (Figure S1) that mainly corresponded to North American populations (1 haplotype), and Eurasian populations (3 haplotypes), as follows: a single main haplotype in North America (with two more minor unique haplotypes in Nanaimo River and Colorado separated by 1 mutational step), one haplotype for most European populations except for most individuals from Spain which had a unique haplotype, and finally a haplotype shared by the individuals in Lake Baikal and one individual in Estonia (Figure S1).

Genetic structuring was evident across the *E. muelleri* SNPs dataset, as shown by STRUCTURE and discriminant principal component analysis (DAPC) (Figure 2). STRUCTURE identified three different optimal scenarios (k = 2, 6, 8), with K = 8 considered optimal based on the Pritchard criterion and K = 2 based on the Evanno criterion (Figure S2). The K2 clustering showed clear differentiation between North American and Eurasian sites (Figure 2A), while the K6 and K8 showed clustering into smaller groups with variable degrees of introgression between them (Figure 2B–C). In the K6 clustering (Figure 2B), within the Canadian sites, Lake O’Connor was entirely assigned to the orange gene cluster, while Nanaimo and Sooke River individuals were predominantly assigned (>90%) to the green cluster, though with slight introgression from the orange cluster. The North American sites were each assigned to a different genetic cluster (Colorado=green, Michigan=blue, and Maine=purple) with Michigan individuals exhibiting genetic admixture from multiple clusters, including blue, orange and purple, and Maine individuals showing introgression from green and blue clusters (Figure 2B). The European (excluding Spain) and Asian samples were predominantly assigned to the red cluster, but showing high levels of introgression from the yellow gene cluster, which fully characterizes the Spanish population at 100% (Figure 2C). In the optimal K8 clustering, groups were very similar, but the substructure within Michigan and Maine was more evident (Figure 2C).

**Figure 2.**
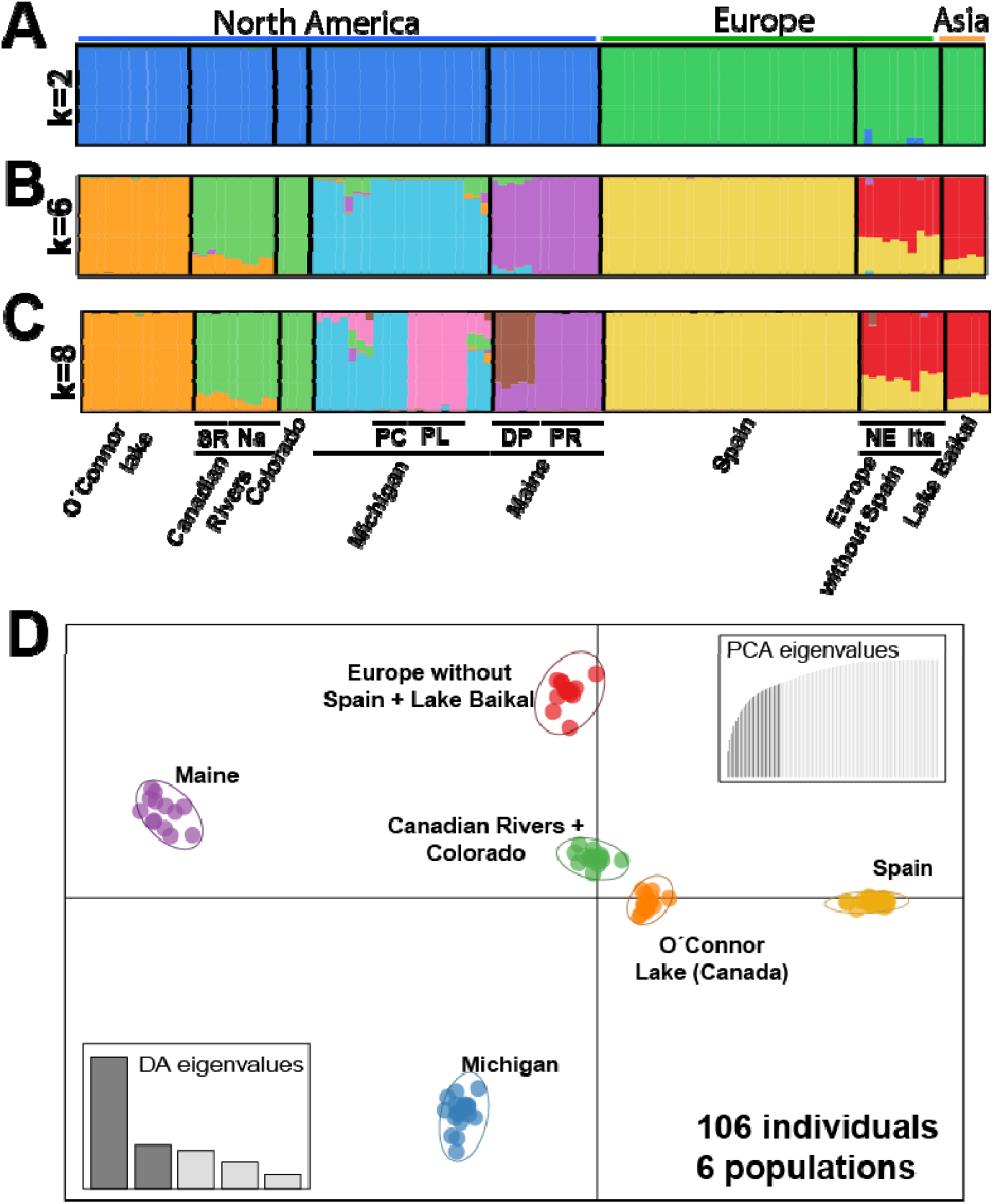
Population structure in *Ephydatia muelleri*. **A**. Admixture plot for K=2. **B**. Admixture plot for K=6. **C**. Admixture plot for K=8. **D**. DAPC analysis of *E. muelleri* showing the samples grouped into six genetic clusters.

Similar to what was observed for the STRUCTURE analysis, the DAPC analysis identified six distinct genetic clusters (k = 6), corresponding to Spain, Europe and Baikal, Lake O’Connor (Canada), Nanaimo and Sooke Rivers (Canada), Colorado, Michigan, and Maine (Figure 2). The separation between populations is mainly reflected along the X-axis, with a notable division between western and eastern Atlantic populations. Additionally, the Y-axis shows clear distinctions between Spain and the Europe/Baikal group.

To understand fine substructuring with the main genetic clusters in *Ephydatia muelleri*, we used fineRADstructure, which revealed a distinct genetic separation between North America and Eurasia (Figure 3A). Within Eurasia, further subdivision was observed, with a clear separation between European and Asian (Lake Baikal) populations, except for the Swedish population, which clustered with the Baikal group. In Europe, samples did not follow a strict geographic pattern and appeared intermixed. In North America, a genetic split was observed between Canadian samples (including those from the Nanaimo and Sooke rivers, and Lake O’Connor) and those from Colorado, Michigan, and Maine. Within these groups, additional subgroups were identified, with each location forming a distinct cluster separate from the others.

**Figure 3.**
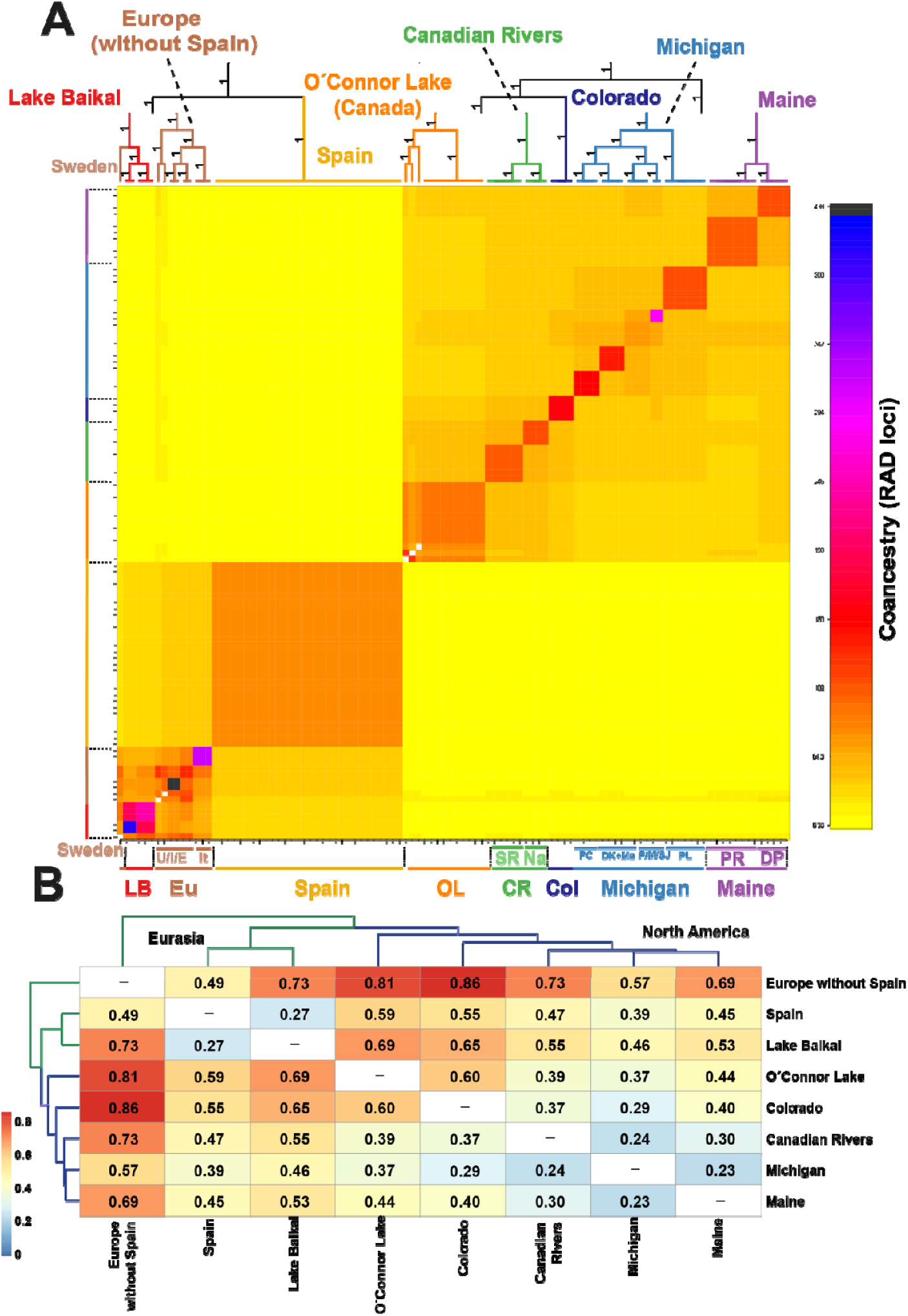
Genetic substructure and isolation. **A**. Coancestry matrix derived from the fineRADstructure analysis. Locations are abbreviated as follows: LB: Lake Baikal, Eu: Europe, Spn: Spain, OL: O’Connor Lake (Canada), CR: Canadian Rivers, Col: Colorado, Na: Nanaimo River, SR: Sooke River, PR: Pemaquid River, DP: Dundee Pond. **B**. Heatmap of F_ST_ values between populations for *E. muelleri*.

Pairwise F_ST_ comparisons among the eight *E. muelleri* groups were all significant, with values ranging from 0.23 to 0.86 (Figure 3B), indicating substantial genetic differentiation between populations. The highest F_ST_ values were observed between Eurasian and North American populations (0.39 to 0.86), indicating greater genetic divergence. Within Eurasia, pairwise F_ST_ comparison of Spain and Baikal showed lower values (0.27) than comparison of Baikal and Europe with Spain excluded (0.73). North American populations generally showed lower pairwise F_ST_ values, with the lowest observed between Michigan and Maine.

Population genetic parameters for the eight *Ephydatia muelleri* sample groups obtained with neutral SNPs are summarized in Table 1. Private allele counts varied widely, with Colorado showing the fewest (29) and Michigan the most (555). Private alleles per individual were lowest in Spain (1.73) and highest in Europe (excluding Spain) at 40.9. The overall expected heterozygosity (*He*) was low (0.108 ± 0.002; mean ± SE), with values ranging from 0.018 ± 0.001 in Spain to 0.105 ± 0.003 in Michigan. Observed heterozygosity (*Ho*) was similarly low (0.050 ± 0.001), ranging from 0.027 ± 0.003 in Spain to 0.080 ± 0.005 in Colorado.

AMOVA results when grouping the samples into two major regions (North America and Eurasia), revealed significant genetic differences between these two groups (Table 2). Significant genetic variation was also observed in the other two comparisons: between regions within continents and between sites within regions. Much of the total variance was attributed to the between-region comparison (28%), although most of the observed variance was attributed to differences between sites.

**Table 2.** Results of the AMOVA analysis for E.muelleri. d.f.: degrees of freedom.Significant p-values in **bold**. The continents represented in this table are North America and Eurasia. The regions include O’Connor Lake (Canada), Nanaimo and Sooke River (Canada), Colorado, the Great Lakes (Michigan), Maine, Spain, Europe (excluding Spain), and Lake Baikal. The sites listed within these regions correspond to the specific locations where each sample was collected.

| Source of variation | d.f. | Sum of squares | Fixation index | % variation | p-value |
| --- | --- | --- | --- | --- | --- |
| Between continents | 1 | 6,74663 | F<sub>CT</sub>: 0.27955 | 27.96 | **0.017** |
| Between regions within continents | 6 | 6,87169 | F<sub>SC</sub>: 0.38777 | 27.94 | **0.000** |
| Between sites within regions | 204 | 15,10013 | F<sub>ST</sub>: 0.55892 | 44.11 | **0.000** |

### 4.2. Demography and Migration Patterns

Our migration analysis using the eight different groupings (O’Connor Lake, Canada; Nanaimo and Sooke River, Canada; Colorado; the Great Lakes, Michigan; Maine; Spain; Europe without Spain; and Lake Baikal), did not detect contemporary migration between North America and Eurasia (Figure 4A). In the western northern hemisphere, Colorado was identified as the present-days source population for the remaining North American populations, with the highest migration rates directed toward Michigan. In addition, migration from O’Connor Lake (Canada) to Michigan was detected. In Eurasia, Spain was identified as the present-days source population for the rest of the populations in this region, with the highest migration rates directed towards the rest of the European countries.

**Figure 4.**
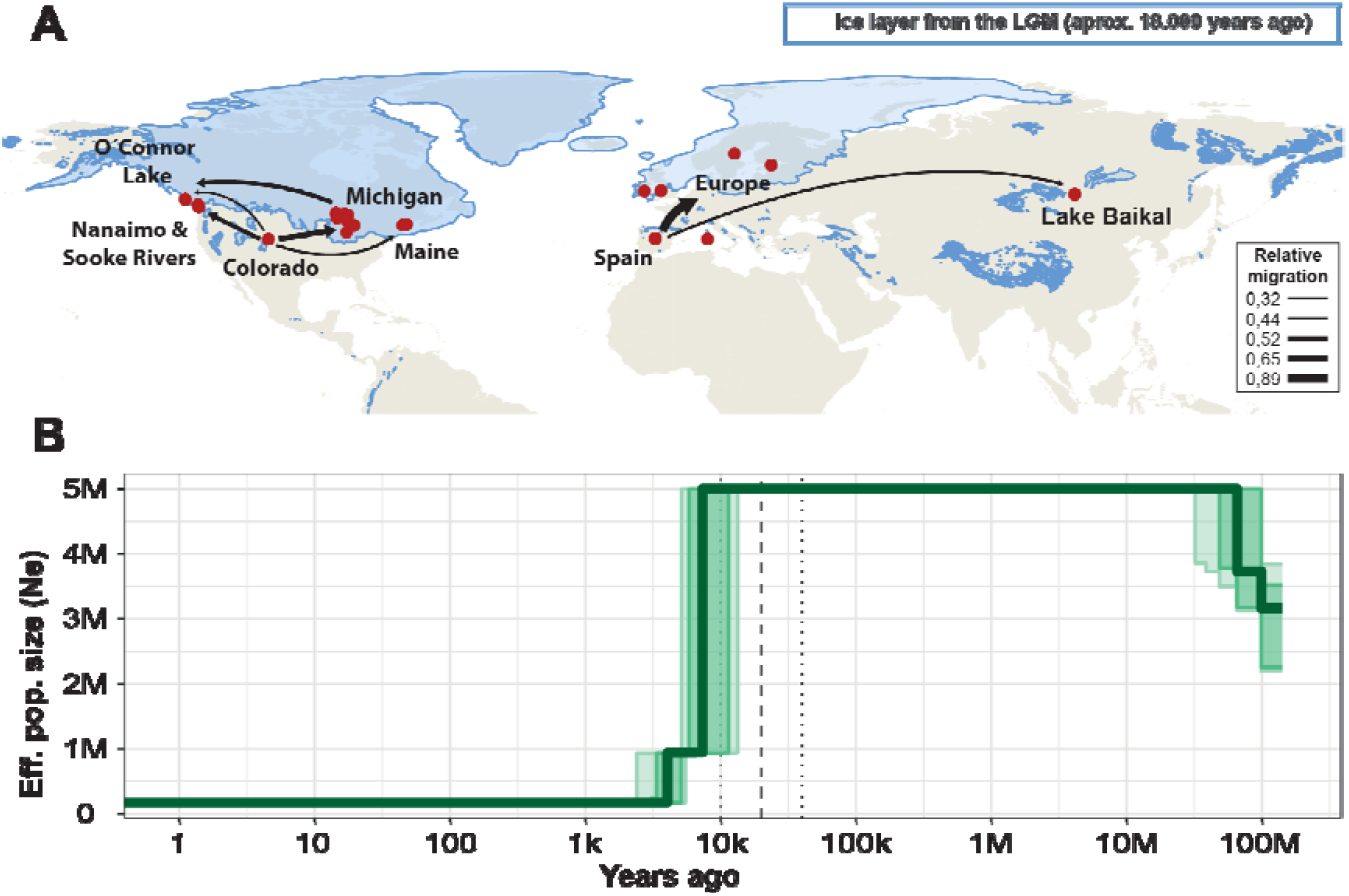
**A**. Observed contemporary migration patterns of *E. muelleri* among the eight geographic areas, as inferred using the divMigrate function of the R package *diverRsity*. The migration map also illustrates areas covered by ice and those that remained ice-free during the Last Glacial Maximum. Data provided by Esri, USGS, FAO, NOAA, USGS, and the University of Koeln. **B**. Demographic history of *Ephydatia muelleri* inferred from ddRADseq data using StairwayPlot2. The bold dashed vertical line indicates the Last Glacial Maximum (20,000 years ago), while the thin dotted lines represent the estimated beginning (40,000 years ago) and end (10,000 years ago) of the Last Glacial Period.

The demographic analysis revealed the dynamics of effective population size (*Ne*) over time, spanning from 100 million years ago to the present. Initially, the population size was notably high, peaking at approximately 5 million individuals in the distant past (Figure 4B). Over time, an abrupt decline in Ne was observed around 10,000 years ago, coinciding with the end of the last glacial period. This trend likely reflects significant historical changes in environmental conditions, genetic diversity, and ecological pressures associated with that era. Following this sharp decline, effective population sizes did not return to its earlier maximum levels.

### 4.2 Population structure and connectivity using SNPs under selection

Of the genome scan (GS) methods used to detect SNPs under selection, BayeScan identified 6 SNPs, Arlequin identified 67, and Baypass identified 73 outlier loci with q-values of < 0.05, producing a set of 139 unique SNPs. Of the GEA methods, pRDA identified 185 potential adaptive SNPs, while LFMM identified 499, for a total of 601 unique SNPs under selection between the two (Figure 5A). In total, all combined methods identified 709 unique loci potentially under selection, with 115 SNPs appearing in at least two different analyses (Figure 5A). Notably, all outlier SNP datasets were found to have relatively even distributions across the genome, with no visible islands of adaptation (Figure 5A, Figure S3).

**Fig 5.**
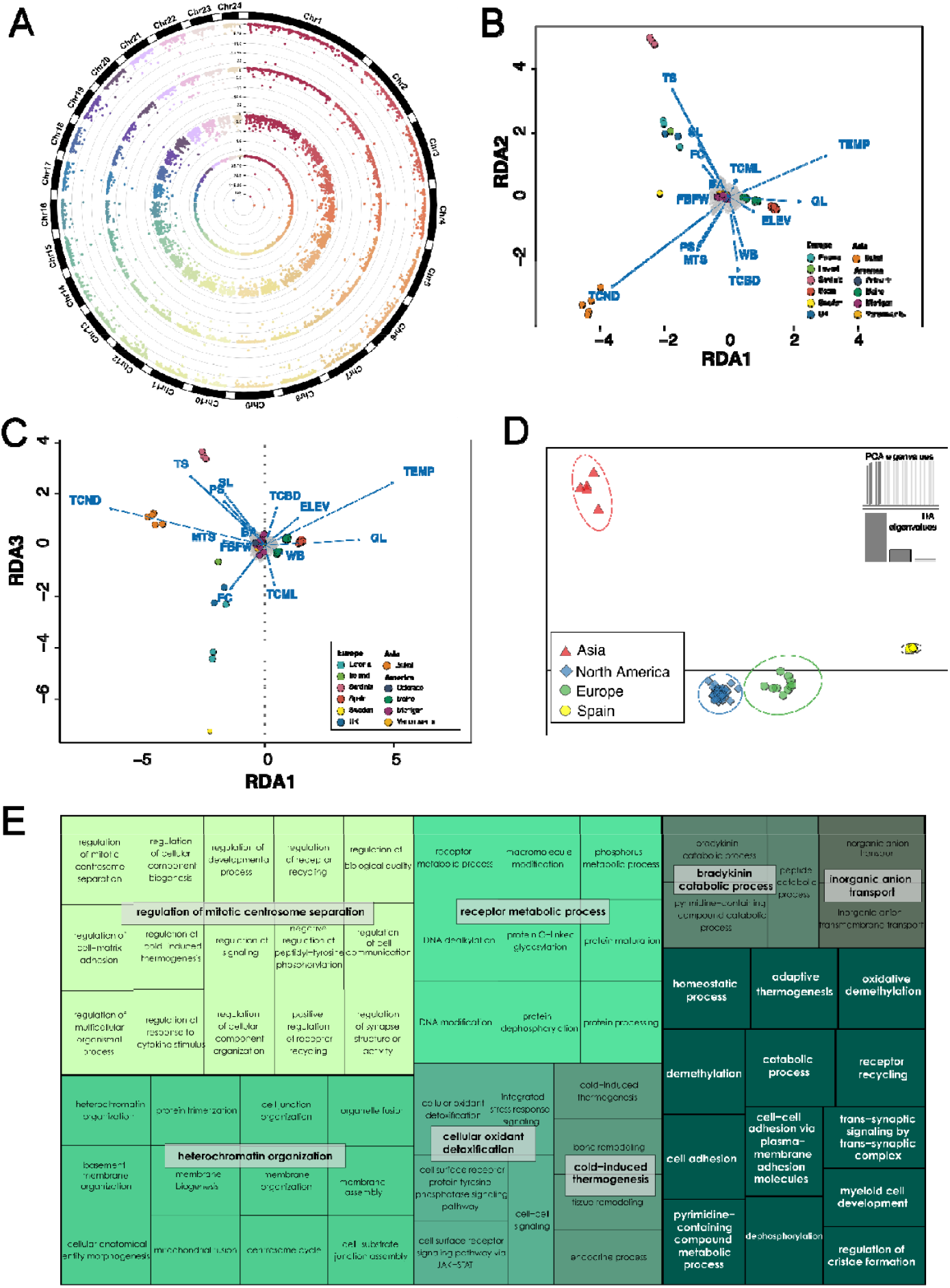
**A**. Manhattan plots of p-values for loci under selection in the analyses: LFMM (PC1), LFMM (PC2), Baypass, Arlequin (in order from the outer circle to the center, respectively). **B-C**. pRDA (axes 1,2 and 1,3, respectively) showing associations of SNP population datasets with environmental factors (Appendix S1), most notably water temperature (TEMP), and tree and plant cover-associated variables (i.e, TS, TCND). **D**. DAPC analysis of putative outlier SNPs, showing groupings of the Americas, Europe, and Lake Baikal. **E**. Treemap showing the enriched Gene Ontologies assigned to the loci under selection that coincided in two of the five adaptive SNP analyses (LFMM, pRDA, Baypass, BayeScan, Arlequin).

The ANOVA of the environmental associations in the pRDA analyses showed that relationships were highly significant (Pr(>F) = 0.001 ***). Additionally, we found R^2^ and adjusted R^2^ values of 0.31 and 0.26, respectively. Outlier SNPs calculated at z = 3 were not evenly distributed across RDA axes; RDA1 yielded only 8 SNPs, while RDA2 yielded 131 and RDA3 yielded 82, with 36 SNPs identified more than once across the axes. While pRDA results did not show clear differentiation of all populations in respective sampling locations, several populations, such as Lake Baikal, Spain, Sardinia, and Maine, were clearly separated along associations with environmental variables (Figure 5B–C). Correlations were also run with the 14 environmental factors and each of the 185 pRDA-identified outlier SNPs, showing that the strongest correlations were represented by only six environmental factors: “tree and shrub cover” (TS), “tree cover, needle-leaved, deciduous” (TCND), fecal coliform concentration (FC), water temperature (TEMP), and grassland cover (GL). Of these, the most common environmental factor was TS across analyses, which was the most highly correlated factor with 117 of the 185 SNPs.

As with the neutral SNP dataset, a DAPC analysis was run on the 115 SNPs identified by at least two outlier SNP analyses (Appendix S2) under the same parameters as mentioned previously. Here, AIC, BIC, and KIC models all grouped the dataset into four clusters (k = 4), demonstrating with high confidence population groupings falling into the geographic categories of North America, Lake Baikal, Spain, and Europe (without Spain) (Figure 5D). We mapped the 115 SNPs to the genome of *E. muelleri*, and recovered 30 different genes, of which only 29 had functional annotations (Table S1). From the genes functionally annotated, we assessed the GO enrichment for relevant Biological Process categories (Figure 5E; Appendix S2), and found: inorganic anion transport (associated to the gene *clcn6*), cellular oxidant detoxification (associated to the gene *pxdn*), cold-induced thermogenesis (associated to the gene *oma1*), bradykinin catabolic process (associated to the gene *ece1*), regulation of mitotic centrosome separation (associated to the gene *kif11*), and heterochromatin organization (associated to the gene *tet1*).

## 5. Discussion

### 5.1. Population structure driven by neutral SNPs in *Ephydatia muelleri*

While there is no established literature on the population dynamics of *E. muelleri*, it is unlikely that any true barriers to dispersal exist apart from the oceans. The hardy, cyclically produced gemmules of freshwater sponges are capable of passive dispersal over large distances by attaching to birds, terrestrial animals, insects, and being picked up by wind (Manconi & Pronzato, 2016), and our globally distributed sampling sites coincide with major avian migration routes, including the Mississippi and American Atlantic Flyways in North America, as well as the East Atlantic and Black Sea/Mediterranean Flyways in Eurasia and northern Africa (Pronzato & Manconi, 1994; Maldonado & Riesgo, 2009; Veen et al., 2005). However, despite the apparent presence of all the necessary tools for dispersal, our findings revealed highly structured populations, with marked genetic differentiation between populations and limited gene flow despite presumably well-connected study sites. This paradoxical genetic structure is highly consistent with the Monopolization Hypothesis (De Meester et al., 2002), which builds on the persistent founder effects hypothesis (Boileau et al., 1992), and predicts high genetic differentiation and limited gene flow across populations of continental aquatic species, passively dispersing cyclical parthenogens due to founder effects amplified by barriers to migration caused by local adaptation and large, resting propagule populations (De Meester et al., 2002).

The first major structure detected in our data was the division in two large, distinct clusters: one corresponding to populations in North America and the other to those in Eurasia. This differentiation is likely a result of large oceanic barriers that restrict gene flow and promote local adaptation, as has been observed in other freshwater species (Lande, 1980; Rahel, 2007). However, beyond the broad genetic divergence between North American and Eurasian populations, our analyses also uncovered marked genetic substructure within each region, with individual populations forming distinct clades that reflect considerable genetic isolation. These findings suggest a pattern that aligns with those observed in other passively dispersed aquatic organisms, such as cladocerans and rotifers (De Meester et al., 2002), which have also been examined under the monopolization hypothesis.

The observed genetic structure, characterized by restricted gene flow even among proximate populations, suggests that historical colonization events have had a more profound influence on shaping large-scale genetic patterns than contemporary dispersal mechanisms. In contrast, current dispersal routes, such as migratory bird pathways, appear to have minimal impact on genetic connectivity. Notably, demographic analysis, along with migration and admixture patterns, reveal a demographic bottleneck event estimated to have occurred approximately 10,000–20,000 years ago, which coincides with significant environmental shifts associated with the retreat of glaciated regions following the Last Glacial Maximum. The impact of this bottleneck on the current population structure is further substantiated by observations in other freshwater invertebrate species, where demographic events such as bottlenecks have been associated with a reduction in effective population size, as well as with elevated F_ST_ values and restricted gene flow in the post-bottleneck phase (Hartl & Clark, 2007; Orsini et al., 2013), a pattern that is similarly evident in our species. These findings suggest that demographic events, such as bottlenecks, play a crucial role in shaping the structural patterns of populations. Specifically, our results showed significant migration from *E. muelleri* populations in Spain to other European sampling sites (Figure 4), which were covered by ice during the Last Glacial Maximum (LGM), alongside pronounced genetic differentiation of the Spanish population across all analyses. This pattern suggests that Spain likely served as a glacial refuge for *E. muelleri* during the LGM. STRUCTURE analyses further indicate the presence of a dominant genotype in Spain, with European and Baikal populations resulting from the admixture of the Spanish genotype with another distinct genetic lineage. These results align with well-established biogeographical patterns of post-glacial expansion from refugia in the Iberian, Italian, and Balkan Peninsulas in Europe (Schmitt, 2007; Taberlet et al., 1998), strongly indicating that modern populations in central and northern Europe emerged from two primary glacial expansion routes—one originating from the Iberian Peninsula and the other potentially from an unsampled refuge in another Southern area. Interestingly, Lake Baikal exhibits migration and admixture patterns similar to those of central European populations, suggesting colonization from European-derived groups. While Lake Baikal was not directly covered by glaciers during the LGM, previous studies have documented the absence of sponge life due to ecological collapse in the lake during this period (Karabanov et al. 2004), thereby supporting the hypothesis that the lake was subsequently colonized by admixed European populations.

In North America, genetic analyses of *E. muelleri* revealed significantly lower F_ST_ values among populations, indicating reduced genetic differentiation and smaller genetic distances compared to those observed in Europe and Asia. This pattern is likely the result of the rapid northward expansion of populations from southern glacial refugia during interglacial periods, as supported by various studies (Hewitt, 2000). Moreover, the south-to-north migration pattern may help explain the genetic flow observed from Colorado to other states, as well as the resulting genetic differentiation between the Colorado population and those in other regions. We also identified a migration pattern originating from Michigan and extending toward Lake O’Connor. This observation aligns with hypotheses proposed in previous studies. For instance, Ding et al. (2017) documented evidence suggesting that populations of quaking aspen migrated from refugia in eastern North America toward the northwest, eventually reaching northern Canada and Alaska as the ice sheets receded following the last glacial period. In light of this evidence, it is plausible that Colorado served as a one of the principal refuges during the LGM. However, the limited number of studies focusing on this region, particularly in comparison to the more extensively studied areas of Eurasia, hinders the ability to draw definitive conclusions. This highlights the need for more comprehensive phylogeographic research on North American species, which would enhance regional understanding and contribute to broader insights into biogeographic patterns (Hewitt, 2000).

An interesting observation is that, apart from the relatively low heterozygosity observed in the populations of this species, in both Colorado and Spain, the presumed founder populations show a notable reduction in genetic diversity and a lower number of private alleles, which is consistent with results from other similar studies on freshwater snails (Redak et al., 2021). This finding contrasts with the typical expectation that founding populations should exhibit higher genetic diversity compared to those in later-colonized areas (De Meester et al., 2002). These observations could suggest two possibilities: first, the loss of genetic diversity may be linked to historical events that caused significant reductions in population size, leading to bottlenecks that reduce heterozygosity and result in allele loss (Hundertmark & Van Daele, 2009). Alternatively, it is possible that the original refugia in these regions (e.g., Southern locations in Central America) have not yet been identified, as our sampling in both Colorado and Spain did not include larger river systems, which may have been critical habitats for maintaining genetic diversity. This pattern has been observed in studies on freshwater snails, where lower genetic diversity is found in populations from tributaries connected to the main rivers (Redak et al., 2021). To determine which of these scenarios is most likely, targeted studies on the history and genetics of populations in both regions are essential. Additionally, sampling in larger aquatic systems would be crucial to explore the potential presence of original refugia.

### 5.2. Genes under selection driving the adaptation in *Ephydatia muelleri*

A key aspect of the monopolization hypothesis is the idea that local adaptation by founder populations serves to limit migration from external populations, further contributing to population differentiation even when geography and physical limitations do not act as barriers to gene flow (De Meester et al., 2002). This is bolstered by evidence that cyclical parthenogens have a heightened capacity for rapid, adaptive evolutionary change through natural selection (Lynch & Gabriel, 1983; De Meester et al., 2002). While a limited number of studies have established evidence of local adaptation in organisms fitting into this framework, such as *Daphnia* (Orsini et al., 2013), none so far have examined the exact mechanisms of adaptation using genome-scale data.

Genomic evidence for local adaptation is critical for verifying the existence of local evolutive processes, a fundamental principle of the monopolization hypothesis, and can shed light on the exact mechanisms of genetic adaptation across small spatial and temporal scales. Importantly, examining the genomic mechanisms at play across populations can also provide insights into whether local adaptation is facilitated by genotypic change or through processes that may not be coded within the genome, such as polyphenism or facultative symbiosis. Despite the established importance of microbial symbiosis in common models of the monopolization hypothesis, such as *Daphnia* (Akbar et al., 2022), symbiotic co-adaptation has been left out of discussion of the eco-evolutive processes involved in the population dynamics of such organisms. Given the evidence for unique genomic adaptations in Spongillidae that have permitted these sponges to cross over into freshwater ecosystems, as well as the large role played by microbial and photosynthetic symbionts in contributing adaptive functions (Manconi & Pronzato, 2016a; Hustus et al., 2023), *E. muelleri* provides an exceptional model through which to investigate the eco-evolutionary processes contributing to local adaptation.

Here, we investigated local adaptations in *E. muelleri* by conducting genome scans and GEA analyses to identify putative SNPs under selection in our sampling sites, as well as potential environmental pressures and biological processes that they may be associated with. These outlier SNP methods yielded a total of 709 putative SNPs under selection, of which 115 coincided with two different adaptive SNP detection methods. Importantly, in addition to a substantial subset of SNPs demonstrating signals of selection, DAPC analysis of these SNPs showed groupings that were clearly distinct from the neutral population structure, instead segregating into the four groups of Lake Baikal, Spain, Europe, and North America, clearly indicating that adaptation is occurring in these populations, likely in response to local selective pressures. Furthermore, the genome-wide distribution of selective SNP supports the segregation of populations indicated in the neutral population structure, as the lack of adaptive islands throughout the genome is consistent with a lack of gene flow that homogenizes linked, non-selective loci and leads to island-shaped adaptive signatures (Tigno & Friesen, 2016). This even distribution of selective SNPs also suggests either that there are many adaptive traits being selected for, or that the adaptive traits are polygenic.

In the pRDA analysis, the adjusted R^2^ value indicated that our environmental factors explained 26% of the variation between populations after accounting for neutral population structure, further supporting the presence of selective pressure by environmental differences across populations. Given the global distribution and high variance in mean annual temperature across our sampling sites, with sample sites including both warm, Mediterranean climate and the Siberian taiga, as well as the resulting implications for differences in annual ice formation, we expected that water temperature would be the most important variable associated with locally adaptive SNPs. However, pRDA revealed that most outlier SNPs detected along the RDA axes were more strongly correlated with local tree and shrub cover, as well as tree type, while the association with temperature, while strong, was consistently weaker.

The association between selected SNPs and water temperature could feasibly be linked with adaptive traits at the gemmule stage, as hatching period has previously been shown to vary according to water temperature (Benfey & Reiswig, 1982). This would provide a clear selective advantage if adapted to local conditions (e.g., gemmules that respond more efficiently to hatching stimuli are more likely to successfully monopolize substrate resources). In the adult stage, physiological adaptations to cold environments are fundamental for this species, because temperatures can vary enormously across the distribution range (Appendix S3), with mean average temperatures of 4.5ºC in Colorado (USA) to 14.3ºC in Sardinia (Italy). Some of the genes identified as under selection by several analyses were in fact involved in processes related to cold-shock and cold adaptation. For instance, *oma1* (*metalloendopeptidase OMA1*) was annotated under the GO category cold-induced thermogenesis (Figure 5E; Appendix S2). Its role in mitochondrial quality control is essential for regulating lipid metabolism as well as for developing the appropriate adaptive response to different metabolic stressors such as a high-fat diet or cold-shock (Quirós et al. 2012). Also, *ece1* (*Endothelin-converting enzyme 1*) appeared under selection in *E. muelleri* (Figure 5E; Appendix S2), which hydrolyzes bradykinin, a molecule elevated in response to acute cold in mice, able to regulate thermogenesis as well (Xiao et al. 2023). The selection of cold-inducible thermogenic factors in sponges is intriguing, since their main role is to increase mitochondrial-based energy expenditure and heat production, largely relying on fat as a fuel source in homeothermic animals (Qiu et al., 2023; Wu et al., 2023), a process that is unknown in poikilotherms (Wu et al., 2023). The role of these thermogenic factors in the physiological adaptations undergone by *E. muelleri* across the Holarctic region is still unknown, but our results open up an exciting avenue for research in this topic.

Besides temperature, other environmental factors were also strongly correlated to the genomic makeup of *E. muelleri*, such as tree and shrub cover, tree cover (needleleaved, deciduous), fecal coliform concentration, and grassland cover. The correlation with tree cover is most likely associated with adaptive responses to the local availability of light and woody substrata, which are essential resources in the life cycle of *E. muelleri*. Previous studies have found preferences for *E. muelleri* in areas with a high abundance of fallen timber, which provides important structural support for colonization of both adults and gemmules, as well as indications that *E. muelleri* can exhibit eco-sensitive phenotypic variations in spicule shape (Dröscher & Waringer, 2007; Andjus et al., 2024).

We also find evidence that *E. muelleri* populations have adapted abilities to more effectively host different local communities of endosymbiotic algae, which play essential roles in providing nutrients to the sponge host (Gost et al., 2023). Previous studies have shown that tree canopy can substantially affect algal communities in rivers and streams by altering the amount of light that penetrates to the water bodies (Roberts et al., 2004; Halliday et al., 2016; Atkinson & Cooper, 2016), and that in the related *Ephydatia fluviatilis*, such differences in light can have dramatic effects on the growth of sponge hosts due to changes in distribution and development of algal symbionts (Gost et al., 2023). Furthermore, previous work has found that microalgal endosymbionts can be transferred vertically from adults to gemmules (Hustus et al., 2023), indicating that endosymbiotic algae are likely an important factor in the founding of populations and subsequent competition between founders and migrants.

The gene-environment associations outlined above are consistent with the monopolization hypothesis if we assume that gemmules with adaptations that improve their fit for hosting local algal symbionts have a selective advantage over gemmules that arrive from water bodies with different light conditions and algal communities, and their corresponding adaptations. However, given the facultatively mutualistic relationship between *E. muelleri* and associated algal communities, and the fact that algae have also been shown to adapt to local light conditions over generations (Roberts et al., 2004), it is also possible that local adaptation is occurring bidirectionally in this system, raising the question of whether the barrier posed to migration by local adaptation, as proposed by the monopolization hypothesis, is at least partially a symptom of coevolutionary adaptation between hosts and symbiont communities in passively dispersing, cyclically parthenogenetic invertebrate holobionts.

Nonetheless, the possibility of local adaptive responses associated with algal endosymbionts is further supported by previous research involving the genes identified to be under selection in these *E. muelleri* populations. Most strikingly, although the *moxd2* and *ubiqp* genes did not yield results from our gene ontology analysis, previous research on transcriptomic responses to algal symbiosis in *E. muelleri* found both of these genes to be upregulated following infection with *Chlorella* symbionts (Geraghty et al., 2021). Specifically, *moxd2* may possibly play a role in cellular-oxidative reduction systems that account for increased ROS following symbiont incorporation, while *ubiqp*, a member of the peroxisome proliferator-activated receptor (PPAR) signalling pathway, is also believed to be involved in the pathway activated by algal infection (Geraghty et al., 2021). Given that previous research has also found differentially regulated responses to introduction of native algal symbionts versus algal symbionts from different geographic areas (Geraghty et al., 2021), our results showing associations between tree cover (i.e., light availability) and adaptive SNPs strongly suggest varied, localized adaptation to algal symbiont communities across the Holartic range of our sampling area.

Similarly, algal symbionts of *E. muelleri* have previously been found to share highly similar microbial communities with their host sponges (Hustus et al., 2023). While there is not yet existing data on the microbial communities of our European and Eurasian sampling sites, previous work on *E. muelleri* sponges sampled from the Sooke River and O’Connor Lake in Canada, as well as Dundee Pond and Pemaquid Rivers in Maine have found overlapping microalgal communities in sponge hosts across these locations (Hustus et al., 2023), as well as overlapping microbial communities of algae and *E. muelleri* hosts, which is consistent with the DAPC results of the SNPs under selection grouping all North American communities in the same cluster. Thus, while future work on the microbial and algal communities of the European and Eurasian populations sampled here is necessary, our results suggest the possibility of genetic adaptations that are responsible for regulating the relationship between *E. muelleri* and algal endosymbionts, or possibly for compensating for variation in local levels of algal photosynthetic activity associated with light availability.

### 5.3 Evidence for Transgenerational Adaptive Responses to Environmental Conditions

Beyond symbiosis-associated genes, we also found signatures of selection on the *ptpra, ptprd*, and *Dicer* genes, which have previously been found to hold a broad diversity of roles in the post-translational regulation of biological processes across different lineages (Byrum et al., 2006; Chen et al., 2017; de Jong et al., 2009; Hu et al., 2023; Iwama & Moran, 2023). The *ptpra* and *ptprd* genes code for protein tyrosine phosphatase receptors, which, in vertebrates, are closely implicated with neuronal function, but more generally are involved with post-translational protein modification and processes such as cell differentiation, adhesion, and growth (Byrum et al., 2006; Chen et al., 2017). *Dicer* genes, on the other hand, have been extensively studied for their importance in RNA interference (RNAi), innate immunity, and epigenetic inheritance (de Jong et al., 2009; Iwama & Moran, 2023). Interestingly, both of these gene groups have been found to be expanded in sponges as compared to other vertebrates, with previous analyses of the *Amphimedon queenslandica* genome showing a highly expanded *ptprd* subfamily in particular, as well as five duplications of the *Dicer* gene, which only has a single copy in vertebrates (Chen et al., 2017; de Jong et al., 2009).

Most interestingly, the *Dicer* gene has been studied extensively across both vertebrate and invertebrate models for its role in regulation of gene expression, and has been demonstrated to act as a developmental switch that facilitates environmental determination of polyphenism in development in *C. elegans* and to play a role in polyphenism in aphid model systems (Grishok et al., 2001; Srinivasan & Brisson, 2012), contributing to transgenerational inheritance of adaptive polyphenism and predictive adaptation in these organisms. This leads us to speculate that *Dicer* may be a key epigenetic mechanism contributing to the phenotypic plasticity of competence for drastically different environments observed in *E. muelleri*. In this scenario, it is possible that *Dicer* may serve as a molecular switch in *E. muelleri*, potentially regulating processes such as the timing of gemmulation, competence for algal infection, or cell growth in response to environmentally determined thresholds.

This hypothesis is particularly compelling considering the importance of light availability for adaptive fit to the local environment in *E. muelleri*, as shown by our results, as well as the high capacity for rapid evolution of organisms with parthenogenetic life phases (De Meester et al., 2002; Lynch & Gabriel, 1983). While no such function has been established in *E. muelleri*, larval settlement and development has been shown to be affected by light conditions in other sponges, which has been speculated to result from polyphenism in development maintained through polymorphisms caused by natural selection on key loci (Degnan & Degnan, 2010). Our results support this proposal, as we detect selection acting on genes demonstrated to be involved in regulation of polyphenic traits in other metazoans (i.e., *Dicer, ptpra, ptprd*). While epigenetics are not mentioned within the framework originally proposed by De Meester (2002), it has nonetheless been observed that epigenetic marks determined by environmental conditions can be passed to the next generation following asexual production in sponges and other invertebrates (Vogt, 2022, 2024). Furthermore, although De Meester et al. (2002) attributes observed local adaptation in *Daphnia* to chance mutations that are amplified by priority effects and rapid clonal evolution under monopolistic habitat colonization, epigenetic principles indicate that environmentally triggered novelties are likely to have a greater evolutionary potential than mutationally-induced ones (Ehrenreich & Pfennig, 2016). Given the preliminary support for selection on genes associated with phenotypic plasticity found here, we propose that local adaptation within the monopolization hypothesis framework may also be attributed to rapid genome evolution induced by genetic assimilation or canalization of environmentally induced polyphenisms under diverging environmental conditions. Such polyphenism in *E. muelleri* may be associated with light and development, although further experimental research is needed to confirm this.

While our gene ontology results for loci under selection in sample *E. muelleri* populations are by no means definitive evidence of polyphenic selection, they nonetheless lend support to future research into this line of investigation. Given that very few studies have established a clear relationship between transgenerational epigenetic marks and environmental fitness, our data suggest that *E. muelleri*, or other aquatic invertebrates that fit within the framework of the monopolization hypothesis, may be a promising model system through which to explore this under-investigated line of research. In particular, the genomic resources available for *E. muelleri*, with its ancestral genes associated with post-transcriptional regulation conserved across metazoans, as well as the wide range of differentiated populations from which to source samples along environmental clines, provide a compelling system through which to explore the role of epigenetic selection and transgenerational modification in climate adaptation. Such research could include experimental studies into *E. muelleri* development, or common garden transplant experiments of different populations, and could potentially provide insights into the role of polyphenism in evolution and speciation, as well as into the future of aquatic ecosystems under climate change.

## 6. Conclusions

We describe the population dynamics of the widely distributed, freshwater sponge *Ephydatia muelleri*, finding evidence of high genetic differentiation between populations and low gene flow despite high dispersal capabilities, with analyses suggesting that large-scale shifts in the environment following the Last Glacial Maximum had more influence on the modern structure of these populations than contemporary dispersal pathways. We also find evidence of local, polygenic adaptation in populations correlating with environmental variables, most notably light availability and water temperature. These findings are highly consistent with the “monopolization hypothesis” (De Meester et al., 2002), suggesting that this may be a viable framework for future research and conservation work involving spongillid species.

However, gene ontology enrichment analysis of our data also indicated that several genes under selection in these populations are associated with gene regulatory processes that have been linked to polyphenic development and transgenerational adaptation in other invertebrates, indicating that epigenetics may play an important role in the evolution and speciation of aquatic invertebrates, which is yet undescribed within the monopolization hypothesis. These novel findings, combined with *E. muelleri*’s wide population distribution, ancestral position within Metazoa, and viability as an easy-to-use model species, suggest that *E. muelleri* may be a promising model for investigating epigenetic processes in metazoans.

