## Supplemental Data 1 for "Genomic connectivity and adaptation signals of the freshwater sponge *Ephydatia muelleri* across its distribution"

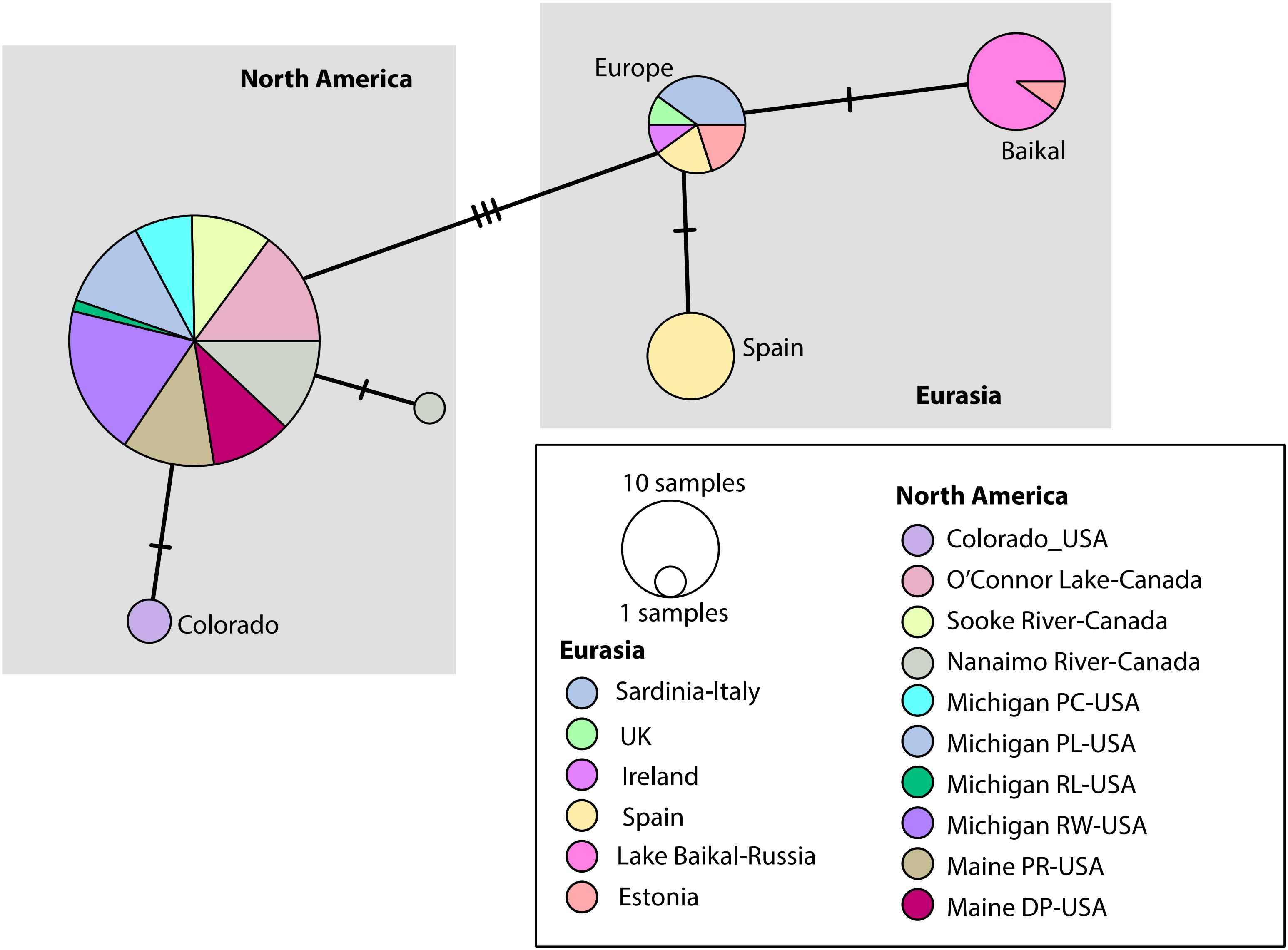

North America
Europe
Baikal
Spain
Eurasia
Colorado
10 samples
1 samples
North America
Colorado_USA
O'Connor Lake-Canada
Sooke River-Canada
Nanaimo River-Canada
Michigan PC-USA
Michigan PL-USA
Michigan RL-USA
Michigan RW-USA
Maine PR-USA
Maine DP-USA
Eurasia
Sardinia-Italy
UK
Ireland
Spain
Lake Baikal-Russia
Estonia

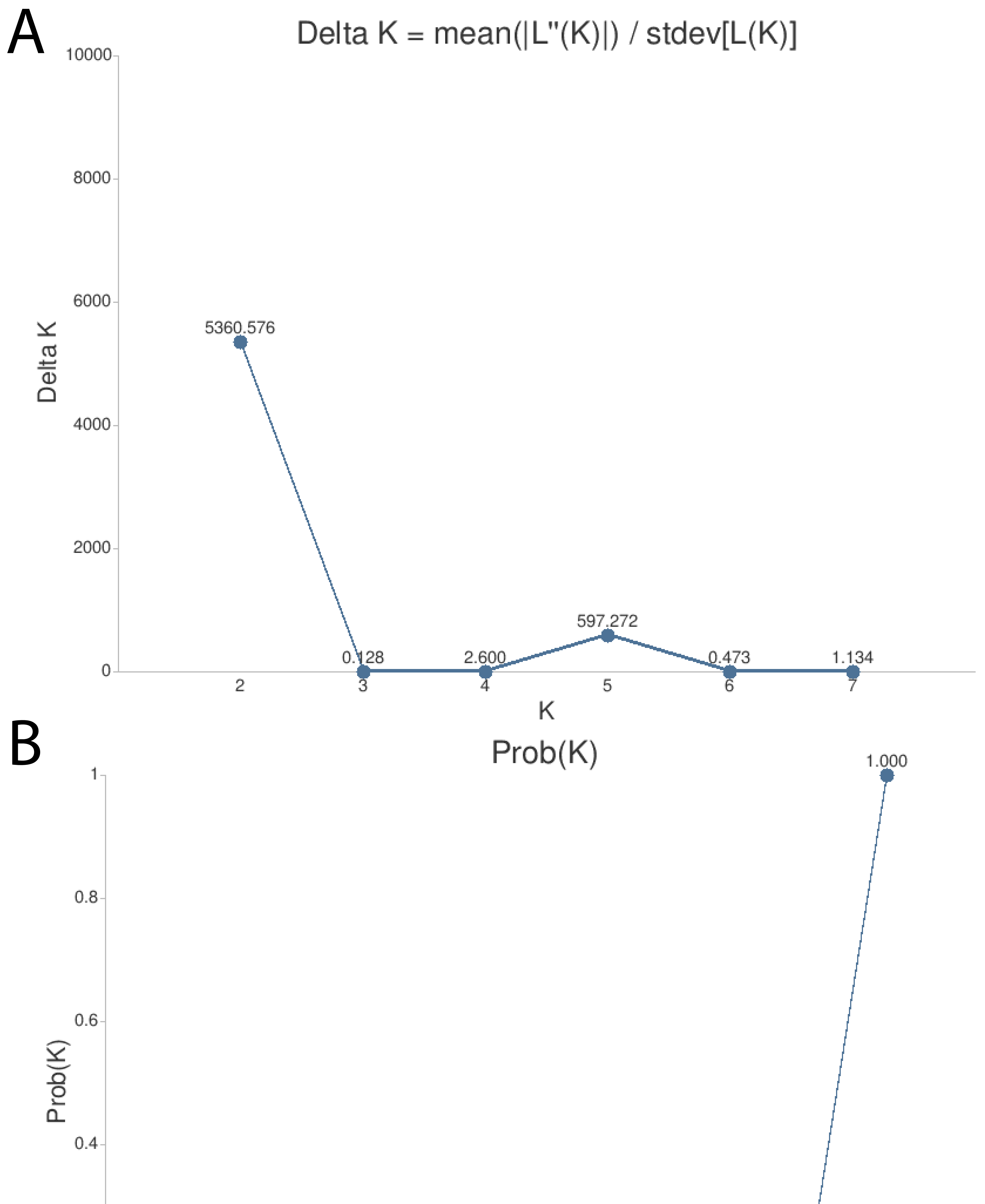

A
Delta K = mean(|L''(K)|) / stdev[L(K)]
Delta K
K
5360.576
0.128
2.600
597.272
0.473
1.134
B
Prob(K)
Prob(K)
1.000

### SUPPLEMENTARY MATERIAL

A

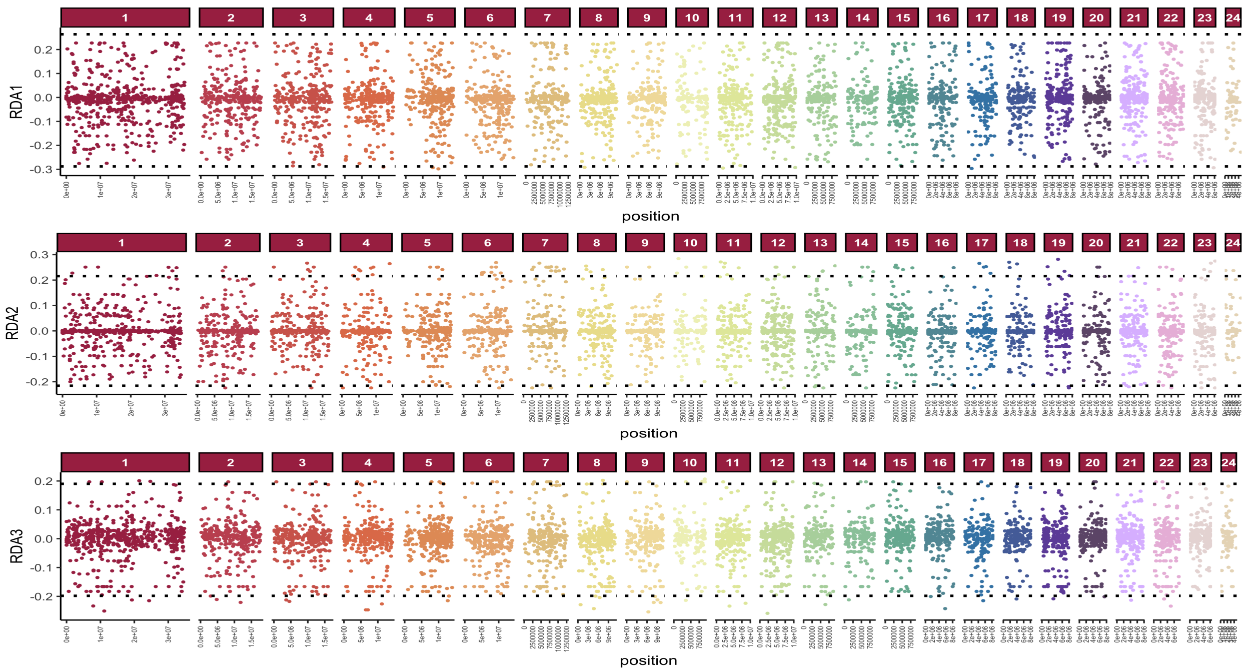


B

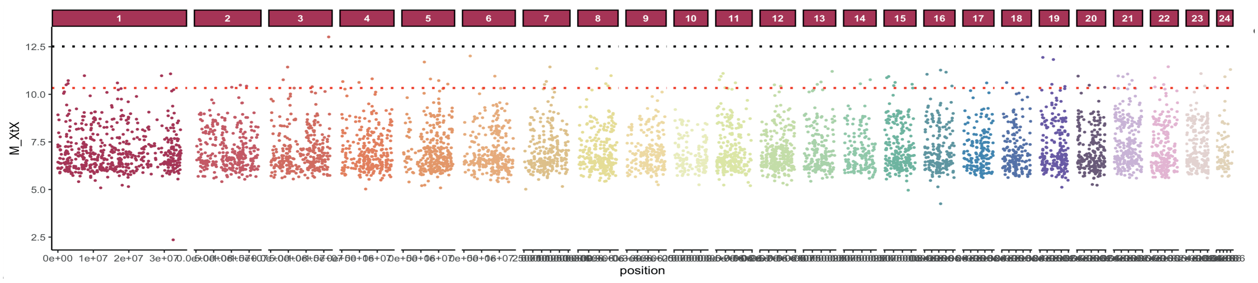


C

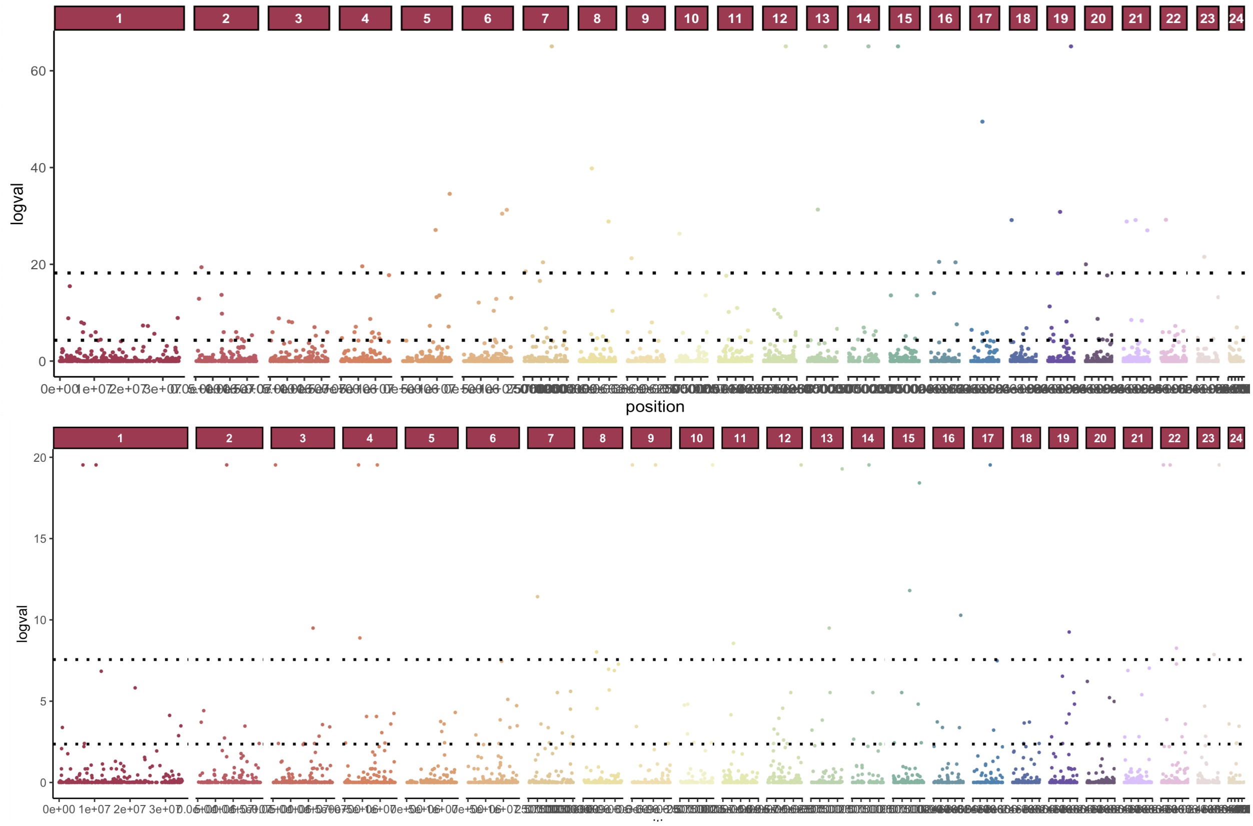


Supplementary Figure S3. Individual Manhattan plots of (A) pRDA z-score outlier axes, (B) Baypass MXtX values, and (C) LFMM -log10 q-values of PCs 1 (above) and 2 (below).
